# Differential involvement of EEG oscillatory components in sameness vs. spatial-relation visual reasoning tasks

**DOI:** 10.1101/2019.12.16.877829

**Authors:** Andrea Alamia, Canhuang Luo, Matthew Ricci, Junkyung Kim, Thomas Serre, Rufin VanRullen

## Abstract

The development of deep convolutional networks (DCNs) has recently led to great successes in computer vision and have become de facto computational models of vision. However, a growing body of work suggests that they exhibit critical limitations beyond image categorization. Here, we study a fundamental limitation of DCNs for judging whether two items are the same or different (SD) compared to a baseline assessment of their spatial relationship (SR). We test the prediction that SD tasks recruit additional cortical mechanisms which underlie critical aspects of visual cognition that are not explained by current computational models. We thus recorded EEG signals from 14 participants engaged in the same tasks as the computational models. Importantly, the two tasks were matched in terms of difficulty by an adaptive psychometric procedure: yet, on top of a modulation of evoked potentials, our results revealed higher activity in the low beta (13-20Hz) band in the SD compared to the SR conditions, which we surmise as reflecting the crucial involvement of recurrent mechanisms sustaining working memory and attention.

**Author Summary:** Despite the impressive progress of deep convolutional networks (DCNs) in object recognition, recent studies demonstrated that state-of-the-art vision algorithms encounter severe limitations when performing certain visual reasoning tasks: for instance, convolutional networks can easily solve problems involving *spatial relations*, but fail in identifying whether two items are identical or different (*same-different task*). This conclusion led us to test the hypothesis that different computational mechanisms are needed to successfully perform these tasks also in the visual system. First, we confirmed in our simulations that DCNs can successfully perform spatial relationship tasks but struggle with same-different ones. Then, we tested 14 participants on the same experimental design while recording their EEG signals. Remarkably, our results revealed a significant difference between the tasks in the occipital brain regions both in evoked potentials and in the oscillatory dynamics. Specifically, an increase of activity was found when performing the SD over the SR condition. We interpret these results as reflecting the fundamental involvement of recurrent mechanisms implementing cognitive functions such as working memory and attention.

## Introduction

The field of artificial vision witnessed an impressive boost in the last few years, driven by the striking results of deep convolutional neural networks (DCNs). Such hierarchical neural networks process information sequentially – through a feedforward cascade of filtering, rectification and normalization operations. The accuracy of these architectures is now approaching – sometimes exceeding – that of human observers on key visual recognition tasks including object [1] and face recognition [2]. These advances suggest that purely feedforward mechanisms suffice to accomplish remarkable results in object categorization, in line with previous experimental studies on humans [3] and animals [4]. However, despite the remarkable accuracy reached in these recognition tasks, the limitations of DCNs are becoming increasingly evident (see Serre, 2019 for a very recent review). Beyond image categorization tasks, DCNs appear to struggle to learn to solve relatively simple visual reasoning tasks otherwise trivial for the human brain [6,7]. A recent study (Kim et al., 2018) thoroughly investigated the ability of DCN architectures to learn to solve various visual reasoning tasks and found an apparent dichotomy between problems: on the one hand stand tasks that require judging the spatial relations between items (Spatial Relationship – SR); on the other, those that require comparing items (Same-Different – SD). Importantly, Kim and colleagues demonstrated that DCNs can learn more easily the first class of problems compared to the second one.

This prompts the question of how biological visual systems handle such tasks so efficiently. Kim et al. (2018) suggest that SR and SD tasks tap into distinct computational mechanisms, thus leading to the prediction that different cortical processes are involved in the two tasks: SR tasks can be successfully solved by feedforward processes, whereas SD tasks seem to require additional computations such as working memory and attention. Here, we tested this compelling hypothesis in two steps: first, we extended Kim’s results by comparing the performance of DCNs on a task in which we directly contrasted SD and SR tasks on the same stimulus set. Second, we recorded electrophysiological responses (EEG) in healthy human participants for the same task, after having matched the difficulty level via an adaptive psychometric procedure. Remarkably, we found that, in addition to a variation in evoked potentials, the SD task elicited higher activity in specific oscillatory components in the occipital-parietal areas, which are typically associated with attention- and memory-related processes. Overall, the present study confirms the computational models’ prediction that the two tasks indeed tap into different computational mechanisms, thus helping to characterize the processes taking place in visual cortex during visual reasoning tasks.

## Results

### Computational modeling

We first extended the results by Kim et al. (2018) for our novel stimulus set: we trained two separate DCN architectures to solve an SD and an SR task using the same stimulus set (Methods). The input to these networks was an image in which two hexominoes were displayed at opposite sides of the screen (see Fig. 1A). The networks were trained to classify whether the two hexominoes were the same or not (SD task) or whether they were aligned more vertically or more horizontally with respect to the midline (SR task).

**Fig.1:**
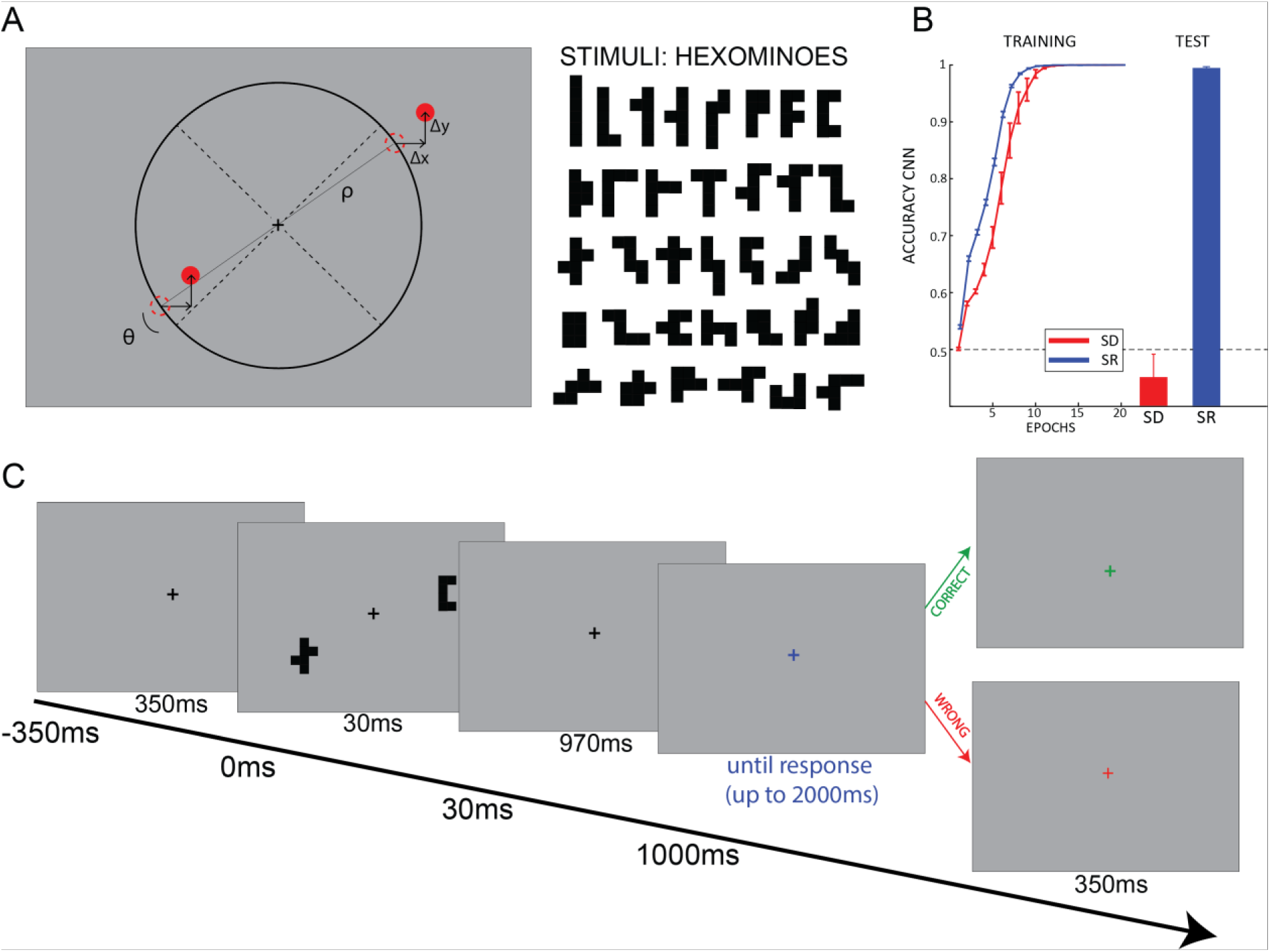
Experimental design and simulation results. A) The items were displayed at opposite sides of the screen (either 45° and 225° or −45° and −225°). Both item positions were jittered by a random amount in both the x and y axis (Δx and Δy in the picture) to make the task non-trivial (i.e. preventing participants from performing the SR task considering only the relative position of one item and its closest diagonal, thus ignoring the spatial relationship between the two items). The items used are hexominoes (right panel). Minimum and maximum item height and width are 1.2° – 3.6° and 1.2° – 2.7° of visual angle, respectively. B) Accuracy of a DCN trained on the same-different (SD; red) and spatial relationship (SR; blue) tasks. The left panel shows the training curves for the two tasks (accuracy over epochs during training); both tasks are fully learnt after 20 training epochs. Test accuracy (right panel), evaluated using novel items never used for training, reveals that the DCN seem to only learn the abstract rule for the SR but not for the SD task, as shown in a previous study. In both panels we show average values ± SE over 10 repetitions using different random initializations. C) At the beginning of each trial a black fixation cross was displayed for 350ms. After 2 stimuli were shown for 30ms, participants waited an additional 970ms before providing the answer. The response was cued by the fixation cross turning blue. After the response, the color of the fixation cross provided feedback: green if the response was correct, red otherwise.

We trained and tested the network on different sets of items (a training and test set, respectively) to assess the networks’ ability to generalize beyond training data. We trained and tested the networks 10 times – randomly initializing networks parameters and training – test set split each time. We report the mean accuracy and standard deviation over these 10 repetitions in Fig. 1B. Our results are consistent with those from Kim et al. (2018): a DCN appears to be able to learn the abstract rule (as measured by the network’s ability to generalize beyond the shapes used for training) for SR tasks much more easily than SD tasks. The effortless ability of humans and other animals [8,9] to learn SD tasks suggest the possible involvement of additional computations that are lacking in DCNs. To test this prediction, we recorded EEG signals from a pool of 14 participants performing the same SD and SR tasks.

### Human behavior

Participants completed 16 blocks using the same stimuli as those used to train DCNs (Fig.1): in half of the blocks they were asked to report whether the two hexominoes were the same or not (SD conditions), in the other half whether the hexominoes were more vertically or horizontally aligned (SR conditions). The two conditions were interleaved in a block design. Participants were required to answer after one second from stimulus onset in order to disentangle motor from visual components in the EEG recordings (Fig. 1C). The QUEST algorithm was used to assure that participants’ accuracy was matched between the two tasks and remained constant throughout the whole experiment. This was done by adjusting two experimental parameters trial by trial (i.e., the hexominoes eccentricity in SD blocks, ρ, and the angle from the diagonal in SR blocks, θ; see Fig. 1A and S1). Maintaining a comparable accuracy between the two tasks reduces the potential for confounds in the electrophysiological analysis due to differences in performance. We confirmed the absence of any substantial behavioral difference between the SD and SR tasks (Fig. 2) with a Bayesian ANOVA on both accuracy (BF_10_ = 0.361, error < 0.001%) and RT (BF_10_ = 0.317, error < 0.89%). In addition, we also investigated each condition separately (Fig. 2, lower panels), comparing the difference between ‘same’ and ‘different’ trials (in SD blocks) and ‘vertical’ and ‘horizontal’ trials (in SR blocks) in both RT and accuracy. All comparisons revealed overall no differences between tasks, except for the accuracy of vertical and horizontal trials in the SR condition, in which the BF proved inconclusive (accuracy: SD - BF_10_ = 0.39, error < 0.012%; SR - BF_10_ = 1.80, error < 0.001%; RT: SD - BF_10_ = 0.333, error < 0.01%; SR - BF_10_ = 0.34, error < 0.01%).

**Fig.2:**
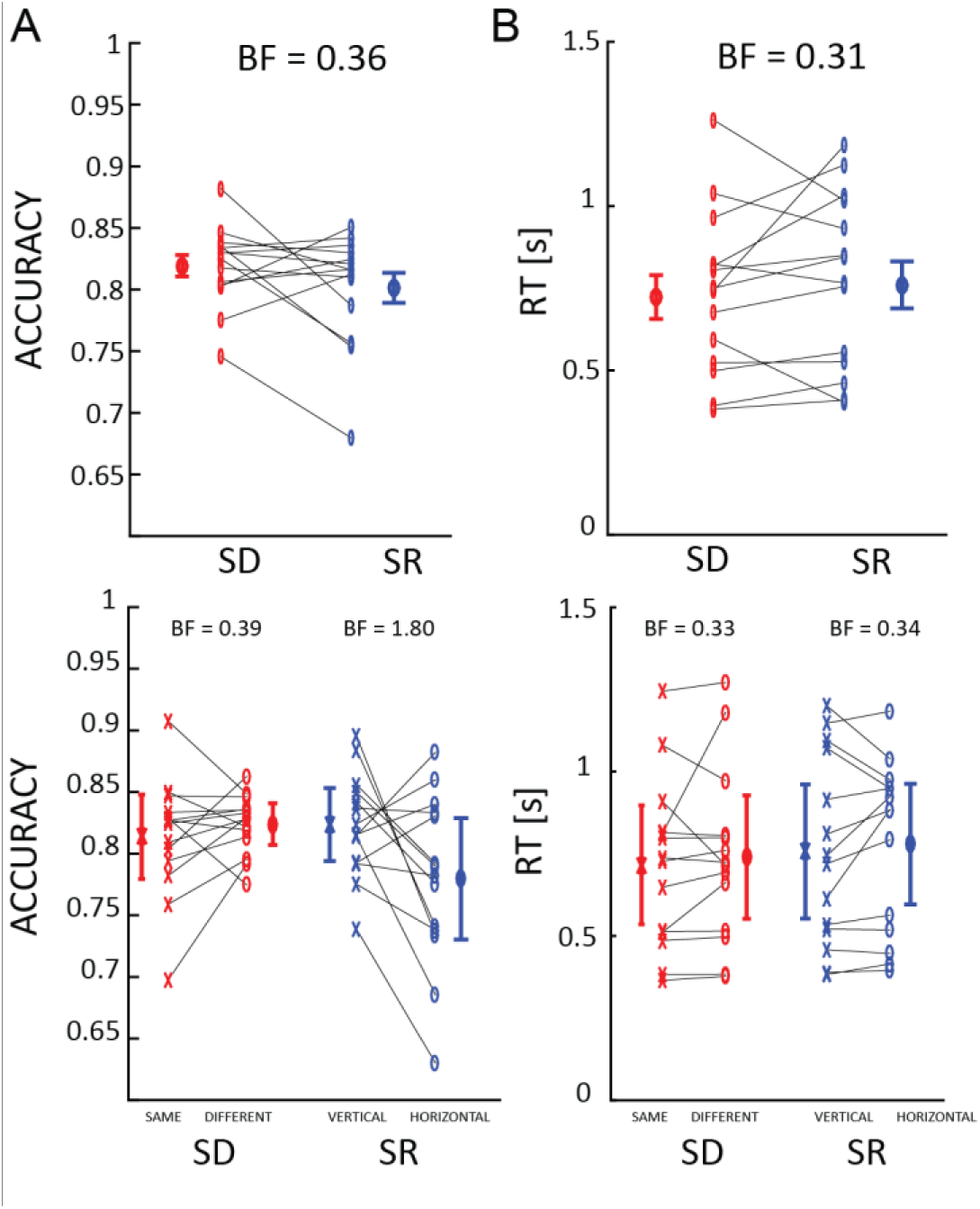
Behavioral results. Humans perform the SD and SR tasks with comparable levels of performance. In the upper panels is shown the average ± SE for accuracy (A) and reaction times (B) for same-different (in red) and spatial relationship (in blue) conditions. Each pair of connected markers represent an individual subject. In the lower panel we show each condition separately (on the x-axis). BF indicates the Bayes factor against the null hypothesis (difference between the two conditions).

### Human electrophysiology: evoked potentials

After having confirmed that performance was equal in the two tasks, we characterized the evoked potentials (EP) in each task. First, we estimated the difference between SR and SD conditions considering 7 midline electrodes (Fig.3, see supplementary Fig. S3 for each condition EPs). The results of a point-by-point t-test corrected for multiple comparisons revealed a significant difference in central and posterior electrodes (mostly Pz and CPz) between 250ms after the onset of the stimuli and the response cue, and the opposite effect in frontal electrodes (FCz and Fz) from 750ms to 1000ms, as confirmed by the topography (Fig.3). Overall these results indicate larger potentials in visual areas during the SD task than in the SR. Previous studies have shown a relation between EP amplitude (particularly P300 and late components) with attention [10,11] and visual working memory [12–14]. Our results are thus consistent with a larger involvement of executive functions in the SD vs. SR task. In the following we investigated whether this hypothesis is corroborated by corresponding oscillatory effects in the time-frequency domain.

**Fig.3:**
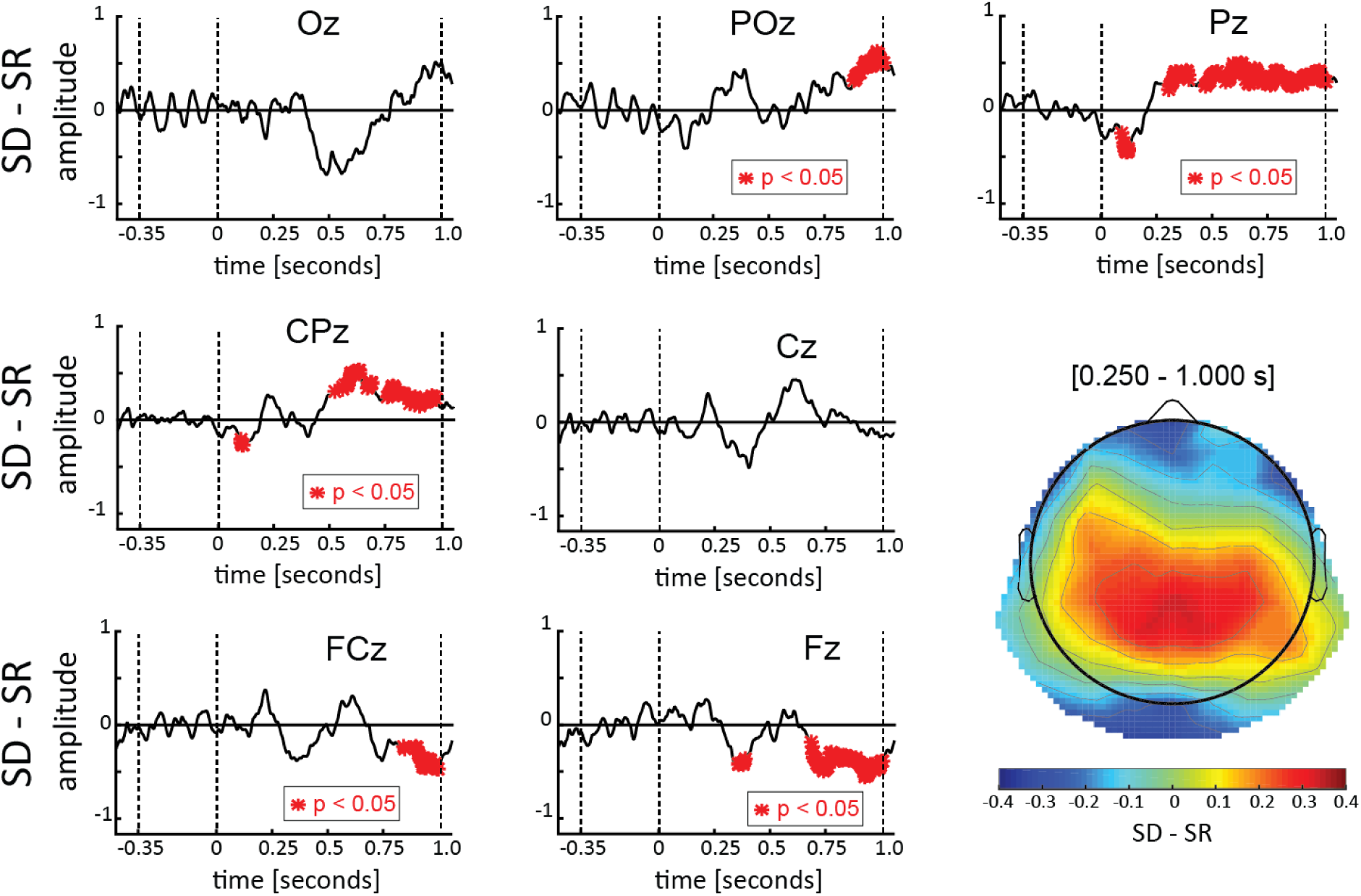
ERPs results. Each panel represents the difference between ERPs elicited in the SD and SR conditions for the 7 midline electrodes. Shown in red are the points for which a difference was found that differs significantly from zero. The results reveal a significant difference from 250ms after stimuli onset until the response cue (at 1000ms) in central parietal regions, and an opposite effect after 750ms in frontal regions. In the bottom-right panel the topography, computed over the 250ms – 1000ms interval, confirmed a larger activity in the SD than in the SR condition (positive difference, warmer colors) in the central-parietal regions, and an opposite effect (negative difference, colder colors) in the frontal regions (and –although not significant– also included occipital regions).

### Human electrophysiology: time-frequency analysis

We performed a time-frequency analysis to try to identify differences between conditions observed in specific frequency bands commonly related to executive functions (e.g., visual working memory). For this purpose, we computed a baseline-corrected log-scale ratio between the two conditions (as shown in Fig. 4A), averaging over all electrodes. Remarkably, a point-by-point 2-tailed t-test corrected with cluster-based permutation test [15] revealed a significantly larger activity in the low beta-band (16-24Hz) in the SD condition between 250 and 950ms after stimuli onset (Fig. 4B). We further quantify the magnitude of the effect by computing the effect size of a one sample t-test against zero averaging per each participant the values within the significant region (t(13)=2.571, p=0.023, Cohen’s d=0.687). The topography of the effect spread mostly over parietal and occipital regions (Fig. 4C), mimicking the topography of the EPs analysis (see supplementary Fig. S4A for the time-frequency plots of each condition separately and corresponding topographies). As previously, these results confirm the prediction that the SD task may involve additional computational mechanisms beyond feedforward computations, possibly indexed by the oscillatory processes identified here. Below, we contextualize and substantiate our results in light of the relevant literature.

**Fig.4:**
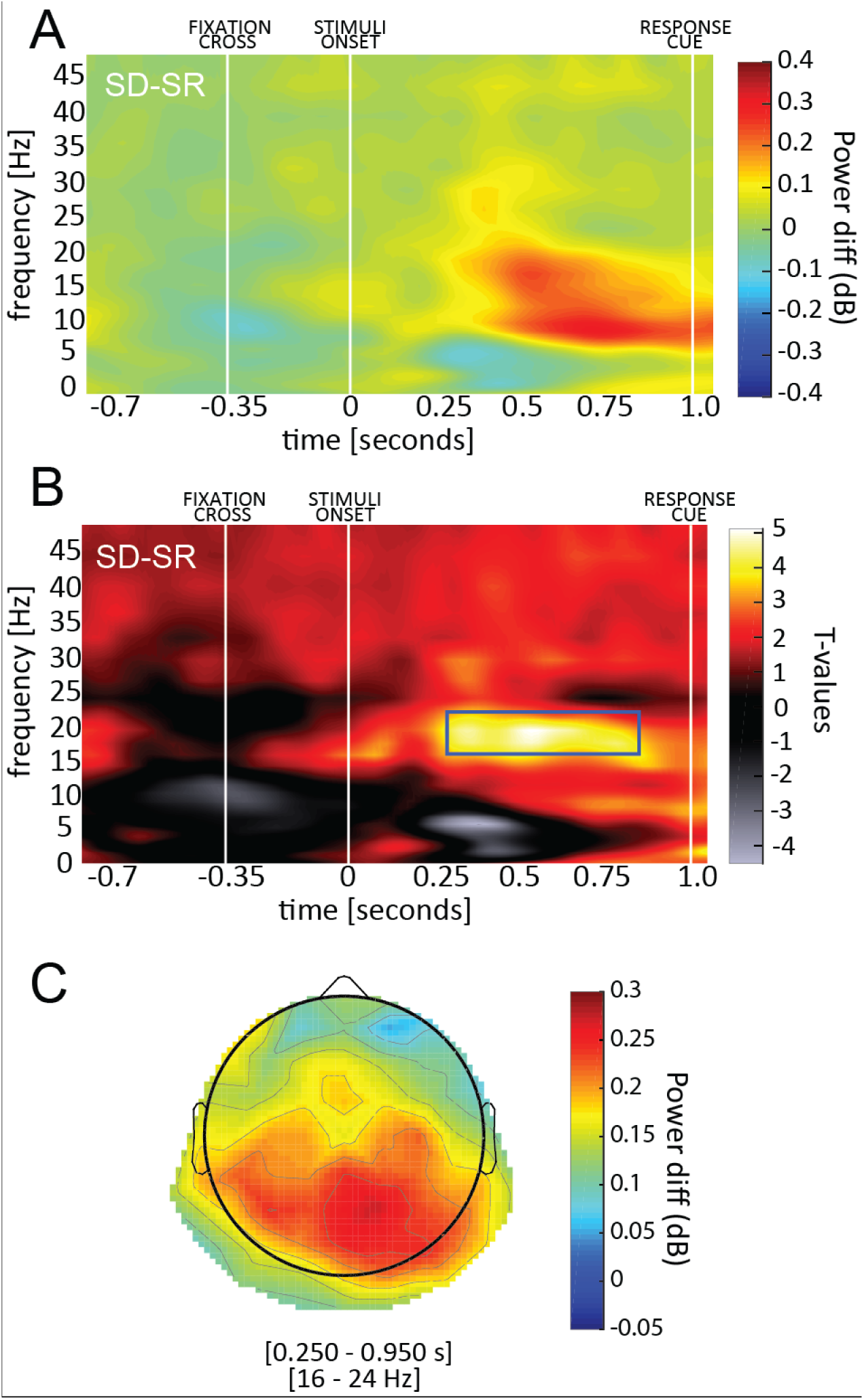
Time-frequency results. A) The difference between SD and SR power spectra is shown in the first panel. White lines indicate the onset of the fixation cross, the stimuli and the response cue. B) The second panel shows the corresponding t values (when testing the difference against zero). We observed a significant region in the low beta band (16-24Hz), between 250ms and 950ms after stimulus onset. C) The topography of the significant time-frequency window reveals the involvement of occipital-parietal regions.

## Discussion

In this study, we confirmed a prediction from the computational study by Kim et al. (2018) that there exists an important dichotomy between visual reasoning tasks: While spatial relation (SR) tasks can be solved by modern deep convolutional networks (DCNs), same-different (SD) tasks pose a significant challenge for these architectures suggesting the need for additional computations beyond a feedforward cascade of filtering, rectification and normalization operations. We have thus investigated the neural strategies used by the human brain to solve SD vs. SR tasks (having the difficulty objectively matched by an adaptive psychometric procedure) and found higher activity in the former in both evoked potentials and oscillatory components. We next interpret these differences as reflecting additional computations carried in the SD task, possibly reflecting working memory and attention processes which are lacking in feedforward architectures such as DCNs.

Computational evidence for the hypothesis that the SD task requires additional computational mechanisms beyond those needed to solve the SR task is provided by Kim et al. (2018). They used a ‘Siamese’ network [16] which processes each stimulus item in a separate (DCN) channel and then passes the processed items to a single classifier network. The authors found that a feedforward network, once endowed with object individuation, could easily learn to solve the SD task. This model simulates the effects of attentional selection, individuation and working memory by segregating the representations of each item. Our results support this interpretation, as revealed by higher activity in the SD compared to the SR task in both evoked potentials and oscillatory frequency bands related to attention and working memory.

Previous work has shown that modulations of beta-band oscillations are related to selective attention mechanisms [17–20]. Different attentional mechanisms may be involved in the two tasks: the SR task could be solved by first grouping items and then determining the orientation of the group [21], whereas the SD task requires the individuation of the two items before comparison. In addition, our results are also consistent with differences in memory processes between the two tasks [22]. One common assumption is that items that are grouped together (as in the SR task) occupy only one working memory slot [23,24], whereas non-grouped items would each hold one slot, resulting in a larger working memory load. Previous literature showed that working memory can also be characterized by neuronal oscillatory signatures. Recent studies have demonstrated an interplay between beta and gamma band frequencies during working memory tasks [25,26]. Similarly, alpha and low beta bands, not only increase with working memory load [27,28], but also in conjunction with the inhibition of competing visual memories in selective memory retrieval [29,30]. All considered, several lines of evidence point towards beta oscillations as crucially involved in both attention and working memory related processes, supporting the conclusion that SD tasks require additional mechanisms compared to SR tasks.

That feedforward neural networks are limited in their ability to solve simple visual reasoning tasks is already being recognized by the computer vision community. Current DCN extensions include modules for integrating local and global features (Chen et al., 2018) as well as recurrent neural architectures (Yang et al., 2018). Our results demonstrate that the human visual system deploys selective attention and working memory to successfully solve visual reasoning tasks: rhythmic cortical oscillations may thus be the signatures of additional computations beyond feedforward processes.

## Materials & Methods

### Participants

Fourteen participants (aged 23–34 years old with a mean age of 27.8 ± 3.1, 5 women, 2 left-handed), volunteered to join the experiment. All subjects reported normal or corrected to normal vision and had no history of epileptic seizures or neurological disorders. The sample size of 14 was chosen based on a previous pilot study performed with the same number of participants on the same task, but without the QUEST adaptive procedure to match the difficulty level between conditions. Importantly, in the previous study we found the very same result (difference between condition in the [16 Hz −24 Hz] ~0.5 dB), with a large effect size (one sample t-test against zero averaging per each participant the values within the significant region, t(13)=7.049, p<0.001, Cohen’s d=1.820). We kept the same number of participants to replicate the effect after having removed the behavioral difference via the QUEST algorithm. This study complies with the guidelines for research at the “Centre de Recherche Cerveau et Cognition” and the protocol was approved by the committee “Comité de protection des Personnes Sud Méditerranée 1” (ethics approval number N° 2016-A01937-44). All participants gave written informed consent before starting the experiment, in accordance with the Declaration of Helsinki, and received monetary compensation for their participation.

### Experimental design

The experiment was composed of 16 experimental blocks of 70 trials each with a total duration of about 1 hour. Each trial lasted ~2 seconds (Fig. 1A): 350ms after the onset of a black fixation cross (0.6° width), 2 shapes were displayed for 30ms on opposite sides of the screen, distant 2*ρ from each other with an angle of ±(45° + θ) with respect to the horizontal midline (ρ being the distance from the center of the screen, and θ the angular difference with the diagonal; see Fig. 1B). Each shape was selected from a subset of 36 hexominoes, a geometric figure composed of 6 contiguous squares (see Fig. 1B) One second after the onset of the hexominoes, the fixation cross turned blue, cuing participants to respond. In half of the blocks, participants had to report whether the two shapes were the same or different (Same-Different – SD condition); in the remaining blocks participants had to judge whether the two stimuli were more aligned horizontally or vertically (Spatial Relation – SR condition). Shapes were displayed at opposite sides of the screen along two main possible orientation axes sampled at random for every trial (either 45° and 225° or −45° and −225°). Both stimuli positions were jittered by a random offset Δx and Δy in both the x and y axis and rotation θ from the main axis. The same offsets were applied to both shapes. The aim of such offsets was to prevent participants in the SR condition from assessing shape orientation by merely comparing their position with an imaginary diagonal line; the offset compels participants to consider the relative position of both hexominoes at once.

Importantly, the difficulty of the two tasks was controlled by an adaptive psychometric procedure (QUEST method, Watson & Pelli, 1983), which varied the eccentricity of the two stimuli ρ (in the SD blocks) or θ (in the SR blocks) to maintain an overall accuracy level of 80% throughout the whole experiment. In fact, larger (smaller) values of ρ made the stimuli more (less) eccentric and the task more (less) difficult; similarly, smaller (larger) values of θ set the stimuli closer to (farther from) the 45° diagonal line, making the task more (less) difficult. We modified one parameter per condition (i.e., per block), while the other was kept constant (using the same value as in the preceding block). After participants responded, they received feedback on their performance: the fixation cross turned green (red) in case of a correct (incorrect) answer. Throughout the experiment the condition blocks were alternated, the first block being the SD condition for all participants. Before starting the first block, participants performed one training block per condition. The purpose of this training was 1) to get participants familiar with the experimental conditions, 2) initialize the ρ and θ parameters in the QUEST method for the first experimental block (initial values were respectively ρ = 5.4° and θ=6°). All experiments were performed on a cathode ray monitor, positioned 57 cm from the subject, with a refresh rate of 160 Hz and a resolution of 1280 × 1024 pixels. The experiment was coded in MATLAB using the Psychophysics Toolbox [34]. The stimuli were presented in black on a gray background. Throughout the experiment we recorded EEG signals.

### EEG recording and pre-processing

We recorded brain activity using a 64-channel active BioSemi electro-encephalography (EEG) system (1,024 Hz digitizing rate, 3 additional ocular electrodes). The pre-processing was performed in MATLAB using the EEGlab toolbox [35]. First, the data was downsampled to 256 Hz. A notch filter [47Hz - 53Hz] was then applied to remove power line artifacts. We applied an average-referencing and removed slow drifts by applying a high-pass filter (>1 Hz). We created the data epochs aligning the data to the onset of the fixation cross. Finally, we performed an ICA decomposition in order to remove components related to eye movements and blink artifacts: we visually inspected the data and removed from 2 to 5 components per subject with a conservative approach (we removed only components in the frontal regions clearly related to eye movements’ activity; Fig. S2 shows the effect of ICA filtering on the V/HEOG in each condition).

### Computational modeling

We extended a previous computational study [7] from which we chose the parameters of the convolutional feedforward network trained on the SD and SR task. Each task was run 10 times, randomly initializing the networks’ parameters and the stimuli used in the training and test set. The network was fed with 150×150 pixel images. Two hexominoes (3×4 pixel size) were placed at opposite side of the screen (see Fig. 1A and ‘Experimental design’). The dictionary of hexominoes was composed of 35 items, which were randomly split between a training (20 items) and a test set (15 items) at each iteration. Both the training and test sets were composed of 1,000 stimuli. The network consisted of 4 convolutional layers. The first layer contained 8 filters of size 4×4. The number of filters doubled at each subsequent layer with a fixed size of 2×2. All convolutional layers used a ReLu activation function with stride of 1 and were followed by pooling layers with 3×3 kernels and a stride of 2. Eventually, three fully connected layers with 256 units preceded a two-dimensional classification layer with a sigmoid activation function. We used binary cross-entropy as a loss function, the Adaptive Moment Estimation (Adam) optimizer [36] and a learning rate of 10e-4. Each simulation was run over 20 epochs with batch size of 50. All simulations were run in TensorFlow [37].

### Statistical analysis – behavior

We analyzed both accuracy and reaction times (RT) by means of Bayesian ANOVA, considering the block condition (SR and SD, see above) as independent variables and the trial condition (whether the stimuli were same or different, or more horizontally or vertically aligned). The result of such analysis provides a Bayes Factors (BF), which quantifies the ratio between statistical models given the data. Throughout the paper, all BFs reported correspond to the probability of the alternative hypothesis over the null hypothesis (indicated as BF_10_). Practically, a large BF (~BF>5) provides evidence in favor of the alternative hypothesis (the larger the BF the stronger the evidence), whereas low BF (~BF<0.5) suggests a lack of effect [38,39]. We performed all Bayesian analyses in JASP [40,41].

### Statistical analysis – electrophysiology

Regarding the EEG recording we performed 2 analyses: one in the time domain measuring Evoked Related Potentials – ERPs, and the other one in the frequency domain using a time-frequency transform. In the first case, we considered the ERPs recorded from 7 midline electrodes (i.e., Oz, POz, Pz, CPz, Cz, FCz and Fz). After subtracting the baseline activity recorded during the 350ms before stimuli onset, we averaged the signals from the SD and SR blocks respectively (i.e., 8 blocks for each condition). Finally, we tested whether the difference between these signals differed from 0 by means of a point-by-point 2-tailed t test with a false discovery rate (FDR) correction for multiple comparisons [42]. Regarding the time-frequency analysis, we computed the power spectra by means of a wavelet transform (1–50 Hz in log-space frequency steps with 1-20 cycles). After baseline correction (i.e., dividing by the averaged activity of the 350ms prior to the onset of the fixation cross), for each participant, we computed the difference in decibel of the two conditions point by point, averaging over all electrodes. As in the ERP analysis, we performed a point-by-point 2-tailed t test to identify the time-frequency regions which were significantly different. We applied a cluster-based permutation to correct for multiple comparisons [15]. First, we identified clusters composed of t values t>3.5 (p<0.01), and for each one we computed the respective global sum. In order to estimate the null distribution over the combined t values, we performed the same procedure 500 times after shuffling the subject by subject SD-SR assignment. Eventually, we obtained the p values for each non-shuffled cluster given the null distribution. All EEG analyses were performed in Matlab; the wavelet transform was performed using the EEGlab toolbox [35].

## Acknowledgments

This work was funded by an ERC Consolidator Grant P-CYCLES number 614244 to RV, a joint CRCNS ANR-NSF Grant “OsciDeep” to RV and TS (IIS-1912280), and two ANITI (Artificial and Natural Intelligence Toulouse Institute) Research Chairs to RV and TS. Additional support to TS was provided by ONR grant (N00014-19-1-2029).

## Supplementary material

**Fig. S1:**
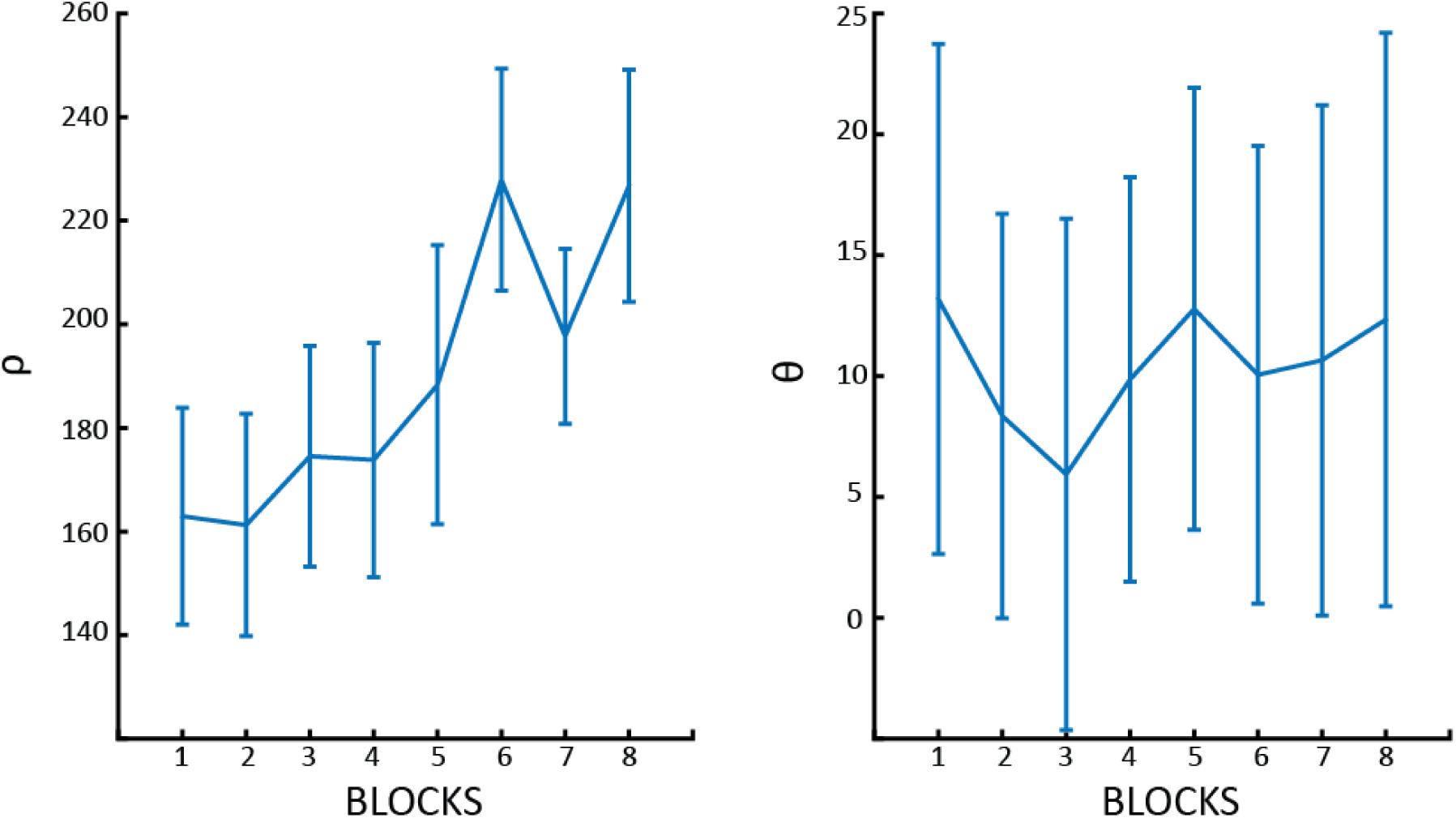
ρ and θ over blocks. Changes over blocks of ρ (the distance between the stimuli - left panel) and θ (the angle between the stimuli and the meridian - right panel) as adjusted by the QUEST algorithm.

**Fig. S2:**
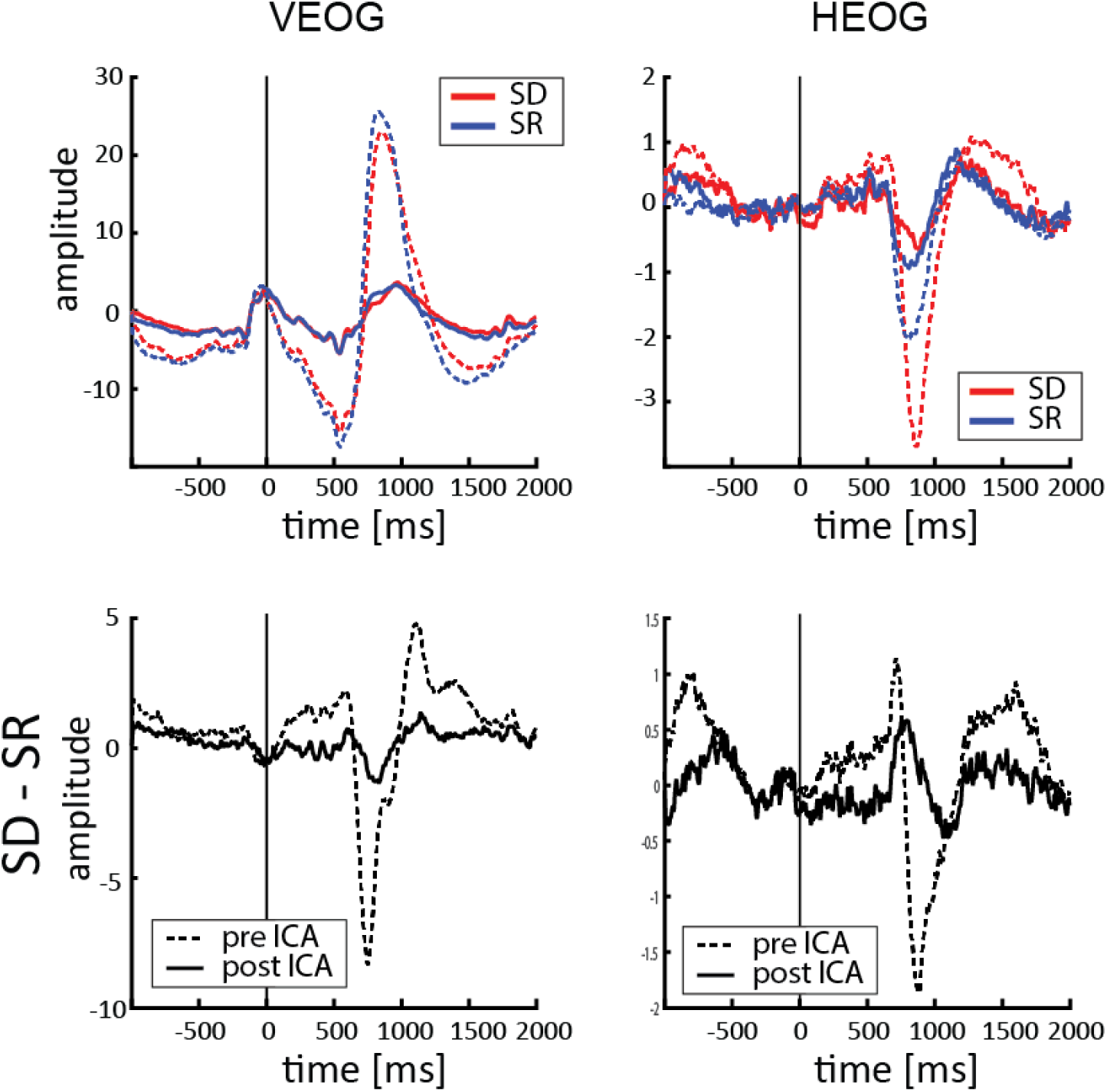
Eye-movement and blink traces. Vertical and Horizontal Electrooculogram before (dashed line) and after (solid line) removing the ICA components related to eye movements. In the top panels both SD (in red) and SR (in blue) conditions are displayed, whereas lower panels show the difference between the two conditions.

**Fig.3S:**
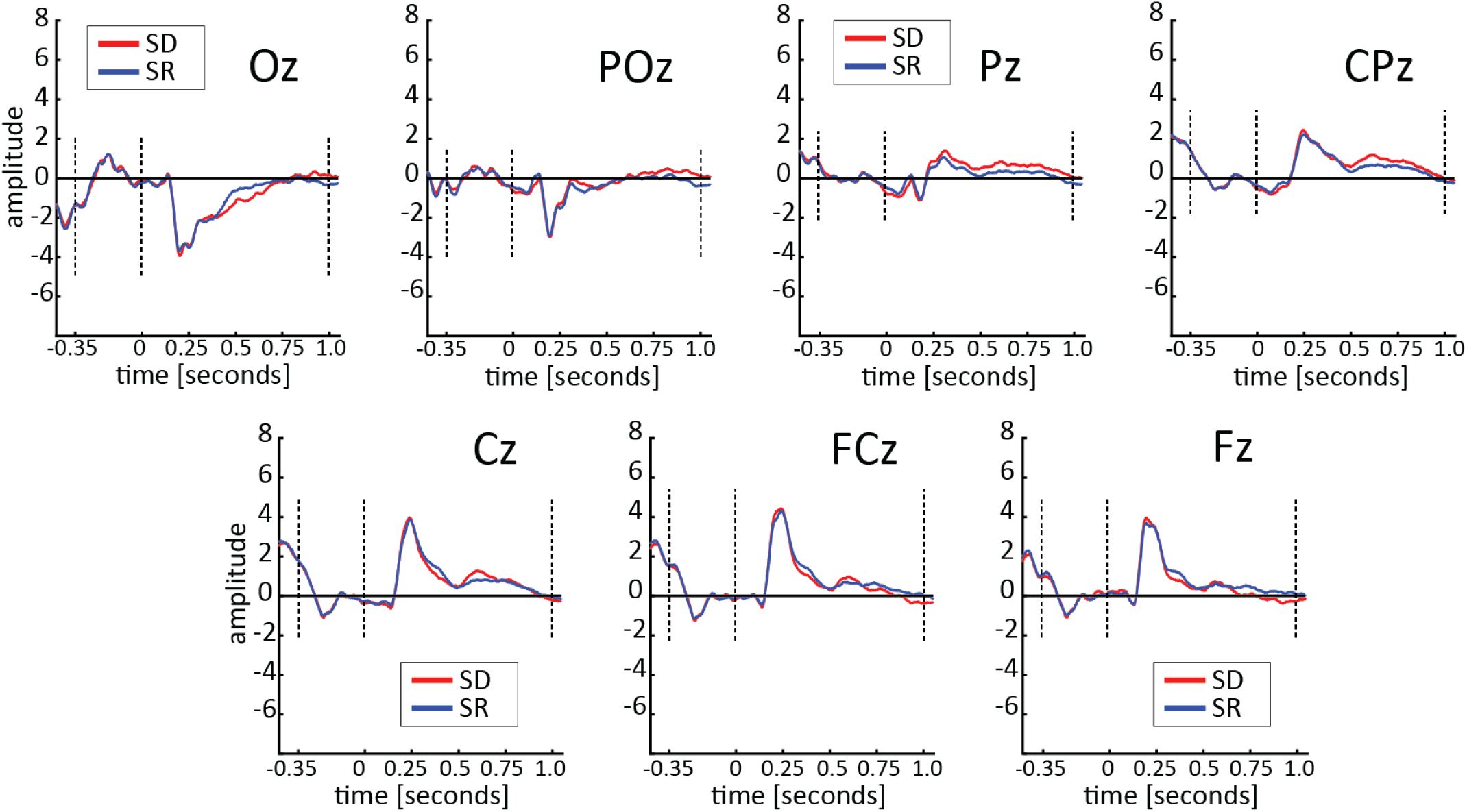
ERPs traces. Baseline-corrected ERPs elicited by the onset of the stimuli (t=0 on the x axis). The two conditions are displayed in red (SD) and blue (SR). Each panel shows one of the 7 median electrodes, from Oz to Fz.

**Fig.4S:**
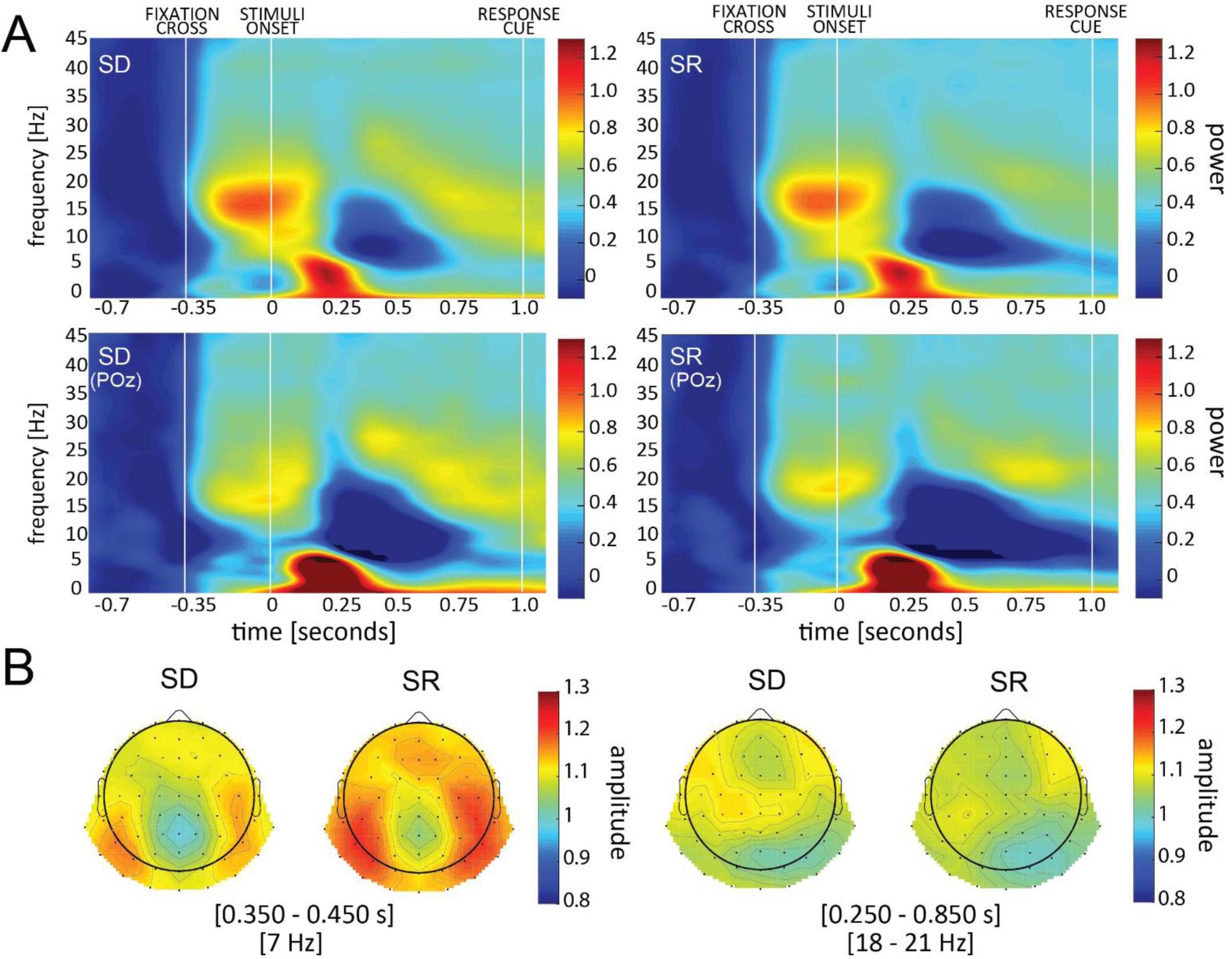
Time-frequency plot. A) Time frequency power baseline corrected (10*log_10_ of the ratio; i.e., units are in decibel) with respect to the 350ms before the onset of the fixation cross. Upper panels reveal the average over all electrodes, lower panel only electrode POz. B) Topography in the time windows where we observed a significant difference between the two conditions (in the theta and beta range respectively).

## References

1. He K, Zhang X, Ren S, Sun J. Deep residual learning for image recognition. Proceedings of the IEEE Computer Society Conference on Computer Vision and Pattern Recognition. 2016. pp. 770–778. doi:10.1109/CVPR.2016.90

2. Phillips PJ, Yates AN, Hu Y, Hahn CA, Noyes E, Jackson K, et al. Face recognition accuracy of forensic examiners, superrecognizers, and face recognition algorithms. Proc Natl Acad Sci U S A. 2018;115: 6171–6176. doi:10.1073/pnas.1721355115

3. VanRullen R, Thorpe SJ. Is it a bird? Is it a plane? Ultra-rapid visual categorisation of natural and artifactual objects. Perception. 2001;30: 655–668. doi:10.1068/p3029

4. Vogels R. Categorization of complex visual images by rhesus monkeys. Part 1: behavioural study. Eur J Neurosci. 1999;11: 1223–1238. Available: http://onlinelibrary.wiley.com/doi/10.1046/j.1460-9568.1999.00530.x/full%5Cnhttp://www.ncbi.nlm.nih.gov/pubmed/10103118

5. Serre T. Deep Learning: The Good, the Bad, and the Ugly. Annu Rev Vis Sci. 2019;5: 399–426. doi:10.1146/annurev-vision-091718-014951

6. Stabinger S, Rodríguez-Sánchez A, Piater J. 25 years of CNNS: Can we compare to human abstraction capabilities? Lecture Notes in Computer Science (including subseries Lecture Notes in Artificial Intelligence and Lecture Notes in Bioinformatics). 2016. pp. 380–387. doi:10.1007/978-3-319-44781-0_45

7. Kim J, Ricci M, Serre T. Not-So-CLEVR: Learning same-different relations strains feedforward neural networks. Interface Focus. 2018;8. doi:10.1098/rsfs.2018.0011

8. Wasserman EA, Castro L, Freeman JH. Same-different categorization in rats. Learn Mem. 2012;19: 142–145. doi:10.1101/lm.025437.111

9. Daniel TA, Wright AA, Katz JS. Abstract-concept learning of difference in pigeons. Anim Cogn. 2015;18: 831–837. doi:10.1007/s10071-015-0849-1

10. Krusemark EA, Kiehl KA, Newman JP. Endogenous attention modulates early selective attention in psychopathy: An ERP investigation. Cogn Affect Behav Neurosci. 2016;16: 779–788. doi:10.3758/s13415-016-0430-7

11. Van Voorhis S, Hillyard SA. Visual evoked potentials and selective attention to points in space. Percept Psychophys. 1977;22: 54–62. doi:10.3758/BF03206080

12. Kok A. On the utility of P3 amplitude as a measure of processing capacity. Psychophysiology. 2001;38: 557–577. doi:10.1017/S0048577201990559

13. McEvoy L. Dynamic cortical networks of verbal and spatial working memory: effects of memory load and task practice. Cereb Cortex. 1998;8: 563–574. doi:10.1093/cercor/8.7.563

14. Fabiani M, Karis D, Donchin E. P300 and Recall in an Incidental Memory Paradigm. Psychophysiology. 1986;23: 298–308. doi:10.1111/j.1469-8986.1986.tb00636.x

15. Maris E, Oostenveld R. Nonparametric statistical testing of EEG- and MEG-data. J Neurosci Methods. 2007;164: 177–190. doi:10.1016/j.jneumeth.2007.03.024

16. Bromley J, Guyon I, Lecun Y, Sickinger E, Shah R, Bell A, et al. Signature Verification using a ‘Siamese’ Time Delay Neural Network. Advances in neural information processing systems. 1994. pp. 737–744. Available: http://papers.nips.cc/paper/769-signature-verification-using-a-siamese-time-delay-neural-network.pdf

17. Richter CG, Coppola R, Bressler SL. Top-down beta oscillatory signaling conveys behavioral context in early visual cortex. Sci Rep. 2018;8. doi:10.1038/s41598-018-25267-1

18. Benchenane K, Tiesinga PH, Battaglia FP. Oscillations in the prefrontal cortex: A gateway to memory and attention. Current Opinion in Neurobiology. 2011. pp. 475–485. doi:10.1016/j.conb.2011.01.004

19. Buschman TJ, Miller EK. Top-down versus bottom-up control of attention in the prefrontal and posterior parietal cortices. Science (80-). 2007;315: 1860–1864. doi:10.1126/science.1138071

20. Lee JH, Whittington MA, Kopell NJ. Top-Down Beta Rhythms Support Selective Attention via Interlaminar Interaction: A Model. PLoS Comput Biol. 2013;9. doi:10.1371/journal.pcbi.1003164

21. Franconeri SL, Scimeca JM, Roth JC, Helseth SA, Kahn LE. Flexible visual processing of spatial relationships. Cognition. 2012;122: 210–227. doi:10.1016/j.cognition.2011.11.002

22. de Fockert JW, G. R, Frith CD, Lavie N. The role of working memory in visual selective attention. S. 2001;291: 1803–1806.

23. Clevenger PE, Hummel JE. Working memory for relations among objects. Attention, Perception, Psychophys. 2014;76: 1933–1953. doi:10.3758/s13414-013-0601-3

24. Franconeri SL, Alvarez GA, Cavanagh P. Flexible cognitive resources: Competitive content maps for attention and memory. Trends in Cognitive Sciences. 2013. pp. 134–141. doi:10.1016/j.tics.2013.01.010

25. Lundqvist M, Rose J, Herman P, Brincat SLL, Buschman TJJ, Miller EKK. Gamma and Beta Bursts Underlie Working Memory. Neuron. 2016;90: 152–164. doi:10.1016/j.neuron.2016.02.028

26. Lundqvist M, Herman P, Warden MR, Brincat SL, Miller EK. Gamma and beta bursts during working memory readout suggest roles in its volitional control. Nat Commun. 2018;9. doi:10.1038/s41467-017-02791-8

27. Pesonen M, Hämäläinen H, Krause CM. Brain oscillatory 4-30 Hz responses during a visual n-back memory task with varying memory load. Brain Res. 2007;1138: 171–177. doi:10.1016/j.brainres.2006.12.076

28. Babiloni C, Babiloni F, Carducci F, Cappa SF, Cincotti F, Del Percio C, et al. Human cortical rhythms during visual delayed choice reaction time tasks: A high-resolution EEG study on normal aging. Behav Brain Res. 2004;153: 261–271. doi:10.1016/j.bbr.2003.12.012

29. Waldhauser GT, Johansson M, Hanslmayr S. Alpha/Beta Oscillations Indicate Inhibition of Interfering Visual Memories. J Neurosci. 2012;32: 1953–1961. doi:10.1523/jneurosci.4201-11.2012

30. Park HD, Min BK, Lee KM. EEG oscillations reflect visual short-term memory processes for the change detection in human faces. Neuroimage. 2010;53: 629–637. doi:10.1016/j.neuroimage.2010.06.057

31. Chen X, Li L-J, Fei-Fei L, Gupta A. Iterative Visual Reasoning Beyond Convolutions. 2018; doi:10.1109/CVPR.2018.00756

32. Yang GR, Ganichev I, Wang XJ, Shlens J, Sussillo D. A dataset and architecture for visual reasoning with a working memory. Lecture Notes in Computer Science (including subseries Lecture Notes in Artificial Intelligence and Lecture Notes in Bioinformatics). 2018. pp. 729–745. doi:10.1007/978-3-030-01249-6_44

33. Watson AB, Pelli DG. Quest: A Bayesian adaptive psychometric method. Percept Psychophys. 1983;33: 113–120. doi:10.3758/BF03202828

34. Brainard DH. The Psychophysics Toolbox. Spat Vis. 1997;10: 433–436. doi:10.1163/156856897X00357

35. Delorme A, Makeig S. EEGLAB: An open source toolbox for analysis of single-trial EEG dynamics including independent component analysis. J Neurosci Methods. 2004;134: 9–21. doi:10.1016/j.jneumeth.2003.10.009

36. Kingma DP, Ba JL. Adam: A method for stochastic gradient descent. ICLR Int Conf Learn Represent. 2015;

37. GoogleResearch. TensorFlow: Large-scale machine learning on heterogeneous systems. Google Res. 2015; doi:10.1207/s15326985ep4001

38. Smith JMB and AFM. Bayesian Theory. Measurement Science and Technology. 2001. pp. 221–222. doi:10.1088/0957-0233/12/2/702

39. Masson MEJ. A tutorial on a practical Bayesian alternative to null-hypothesis significance testing. 2011; 679–690. doi:10.3758/s13428-010-0049-5

40. JASP Team. JASP (Version 0.8.6.0) [Internet]. [Computer software]. 2018. Available: http://jasp-stats.org

41. Love J, Selker R, Verhagen J, Marsman M, Gronau QF, Jamil T, et al. Software to sharpen your stats. APS Obs. 2015;28: 27–29.

42. Hochberg B. Controlling the False Discovery Rate: a Practical and Powerful Approach to Multiple Testing. J R Stat Soc. 1995;57: 289–300. doi:10.2307/2346101

